# Nuclear Export Through Nuclear Envelope Remodeling in *Saccharomyces cerevisiae*

**DOI:** 10.1101/224055

**Authors:** Baojin Ding, Anne M. Mirza, James Ashley, Vivian Budnik, Mary Munson

## Abstract

In eukaryotes, subsets of exported mRNAs are organized into large ribonucleoprotein (megaRNP) granules. How megaRNPs exit the nucleus is unclear, as their diameters are much larger than the nuclear pore complex (NPC) central channel. We previously identified a non-canonical nuclear export mechanism in *Drosophila* (Speese et al., *Cell* 2012) and mammals (Ding et al., in preparation), in which megaRNPs exit the nucleus by budding across nuclear envelope (NE) membranes. Here, we present evidence for a similar pathway in the nucleus of the budding yeast S. *cerevisiae*, which contain morphologically similar granules bearing mRNAs. Wild-type yeast displayed these granules at very low frequency, but this frequency was dramatically increased when the non-essential NPC protein Nup116 was deleted. These granules were not artifacts of defective NPCs; a mutation in the exportin *XPO1* (CRM1), in which NPCs are normal, induced similar megaRNP upregulation. We hypothesize that a non-canonical nuclear export pathway, analogous to those observed in *Drosophila* and in mammalian cells, exists in yeast, and that this pathway is upregulated for use when NPCs or nuclear export are impaired.

**SUMMARY:** Ding et al., describe a non-canonical mRNA export pathway in budding yeast similar to that observed in *Drosophila*. This pathway appears upregulated when the NPC is impaired, nuclear envelope integrity is disrupted, or the export factor Xpo1 (CRM1) is defective.

## INTRODUCTION

A fundamental dogma in eukaryotic cell biology is that trafficking of endogenous cellular materials into and out of nuclei occurs exclusively through the nuclear pore complex (NPC) (Aitchison and Rout, 2012; Grunwald et al., 2011). However, in many eukaryotes, subsets of exported mRNAs are organized into large ribonucleoprotein (RNP) particles containing not only mRNAs, but also regulatory proteins that control their translation at precise cellular sites and in response to specific stimuli (Martin and Ephrussi, 2009). A major issue is how these large complexes (~100-700 nm) can exit the nucleus through NPCs, as the diameter of the NPC channel is ~40-50 nm (Musser and Grunwald, 2016)). A prevailing model proposes that very large ribonucleoprotein (RNP) particles are not assembled in the nucleus, but rather that the components are individually transported out through NPC, and assembled in the cytoplasm (Moore, 2005). However, an alternative mechanism of nuclear export, affected by remodeling nuclear envelope (NE) membranes, has been demonstrated for viruses of the Herpes type (HV; (Hellberg et al., 2016)). In this pathway, capsids packaging viral DNA are assembled in the nucleus, but are much too large to escape the nucleus through NPCs. Therefore, HV use a three-step process to exit the nucleus through the NE. In the first step, viral and cellular kinases disrupt the thick nuclear lamin structure by phosphorylating lamina components, thus gaining access to the inner nuclear membrane (INM). In the second step of “primary envelopment,” the capsid reorganizes the INM to become enveloped by this membrane, which is then pinched off and released into the perinuclear space. Finally, the membrane surrounding the capsid fuses with the outer nuclear membrane (ONM) in a process of “de-envelopment”, releasing a naked virion that undergoes secondary envelopment and maturation in the cytoplasm. While in principle this mechanism does not require NPCs, whether NPC components might be involved in any aspect of this mechanism is unknown. Substantial progress has been made in identifying viral and cellular proteins capable of carrying out this process during nuclear HV egress (Darlington and Moss, 1968; Hagen et al., 2015; Hellberg et al., 2016; Lee and Chen, 2010; Lye et al., 2017; Mettenleiter et al., 2013; Roller, 2008). However, a long-standing question is whether NE-budding is exclusive to the HV life cycle, or is endogenously used by eukaryotic cells for normal cellular processes.

Previously, a non-canonical mRNA nuclear export mechanism was reported, which resembled that of HV at the *Drosophila* neuromuscular junction (NMJ) during synaptic growth (Speese et al., 2012). This process was also observed in mammalian neurons during differentiation (Ding et al, in preparation). In this pathway, mRNAs are assembled into megaRNP granules in the nucleus, and are transported into the cytoplasm, likely by remodeling NE membranes in a manner similar to HV. A key component in the *Drosophila* and mammalian megaRNP nuclear egress pathway is the AAA+ ATPase, Torsin or TorsinA (TorA) respectively. While 4 *torsin* paralog genes are encoded in the mammalian genome, a single *torsin* gene in encoded by the *Drosophila* genome, circumventing complications in the analysis of single mutants arising from compensation. In humans, mutations in TorA cause DYT1, a severe form of dystonia and this is associated with defects in the ER and NE blebbing (Naismith et al., 2004). In *Drosophila*, either expression of a mutant torsin modelled after the DYT1 mutation, eliminating, or downregulating Torsin, leads to accumulation of megaRNP granules that bud into the perinuclear space, but remain tethered to the INM (Jokhi et al., 2013).

The components of the canonical RNA and protein import and export pathways through the NPC have been well characterized (Kohler and Hurt, 2007; Natalizio and Wente, 2013). The major export pathways employ the exportins CRM1 (Xpo1 in S. *cerevisiae)* and TAP (Mex67). In contrast, the mechanisms of the structural and regulatory machinery of the endogenous megaRNP export pathway are mostly unknown (Jokhi et al., 2013; Speese et al., 2012), despite progress in identifying the viral machinery (Funk et al., 2015; Hagen et al., 2015; Liu et al., 2015; Maric et al., 2014; Maric et al., 2011; Turner et al., 2015). Additional functional questions remain: what are the variety and extent of the cargos carried by the endogenous pathway? Is this pathway used for transport of specific mRNAs that are packaged together for local translation, as predicted for megaRNPs at the NMJ? Is this pathway used for clearance of large protein aggregates and defective NPCs (e.g., (Laudermilch and Schlieker, 2016; Webster et al., 2016))? Is this pathway conserved across different eukaryotic species, and how did the pathway evolve? What are the cellular processes during which the pathway is used (e.g. during *Drosophila* NMJ development (Fradkin and Budnik, 2016; Speese et al., 2012), or proper mitochondrial function (Li et al., 2016)?

To begin addressing such questions, we sought to determine if this pathway is also present in the biochemically and genetically tractable model organism, the budding yeast S. *cerevisiae*. Previously, mutations in several different yeast genes were observed to result in abnormal NE morphology, including the observation of electron-dense vesicular granules at the perinuclear space, reminiscent of the megaRNP granules observed in *Drosophila* and mammals. Similar granules were detected in various yeast mutant strains, including deletions or mutations of the nucleoporins Nup116 (Wente and Blobel, 1993), Nup145 (Teixeira et al., 1997; Wente and Blobel, 1994) and Nup188 (Nehrbass et al., 1996); the acetyl Coenzyme A carboxylase protein Acc1 (Schneiter et al., 1996); the nuclear envelope proteins Apq12 (Scarcelli et al., 2007) and Brr6 (de Bruyn Kops and Guthrie, 2001; Hodge et al., 2010); and the nuclear-associated p97 co-factor Npl4 (DeHoratius and Silver, 1996). In several of these studies, it was postulated that the nuclear envelope defects and granules were an aberrant accumulation of material trapped due to abnormal blockage of transport when NPCs were defective (e.g., (Wente and Blobel, 1993)). For other mutants, the nuclear envelope membrane integrity appeared disrupted (Hodge et al., 2010; Schneiter et al., 1996). We hypothesized that, as in other species, these granules could represent a nuclear transport mechanism in budding yeast for ultra-large RNPs, which functions as a parallel or backup physiological mRNA and protein export pathway.

## RESULTS AND DISCUSSION

### megaRNP-like granules are present in wild-type yeast cells

To investigate the presence of the non-canonical NE export pathway in S. *cerevisiae*, we examined the wild-type (WT) strain W303 using thin section transmission electron microscopy (TEM) of yeast fixed by high pressure freezing (HPF). We discovered that WT cells contain electron dense granules at the perinuclear space, between the INM and the ONM (Fig. 1A-E; RNP, black arrows), albeit at a very low frequency (~ 1%) at 37 °C. In contrast, these granules were seldom observed in WT cells at 25°C. The slight increase in granule frequency at the elevated growth temperature (37 °C) may correspond to upregulation of transcription in response to heat shock (McAlister et al., 1979; Miller et al., 1979). The granules, always found as singlets localized within the perinuclear space (Fig. 1A1-2), appeared surrounded by membrane (e.g., Fig. 1C2; white arrow), had some internal variability in electron density (Fig. 1A2; 1C2), and ranged from 60-150 nm in diameter (Fig. 1F). In addition, while most of these granules were close to, but with variable degrees of apposition to the INM, they were usually detached from the ONM (Fig. 1 C2; 1F). The region of apposition between the granules and INM appeared much wider than NPCs (see Fig.1A1, white arrowhead for an NPC cross-section, grey arrowheads for NPC sagittal sections [shown at high magnification in A2] and Fig.1F for quantification). At regions of the NE containing a granule, the ONM was considerably separated from the INM, forming an enlarged perinuclear space Fig. 1A2; 1F). In contrast, NPCs (Fig. 1A1; white arrowhead) and spindle pole bodies (SPB; Fig.1D; black arrowhead) were observed only in regions of the NE outside of these enlargements, and the perinuclear space around NPCs or SPBs was much smaller and of consistent size (Fig.1F). Granules appeared smaller (93 ± 7 nm in diameter in yeast compared to ~200 nm in flies (Speese et al., 2012)).

**Figure 1.**
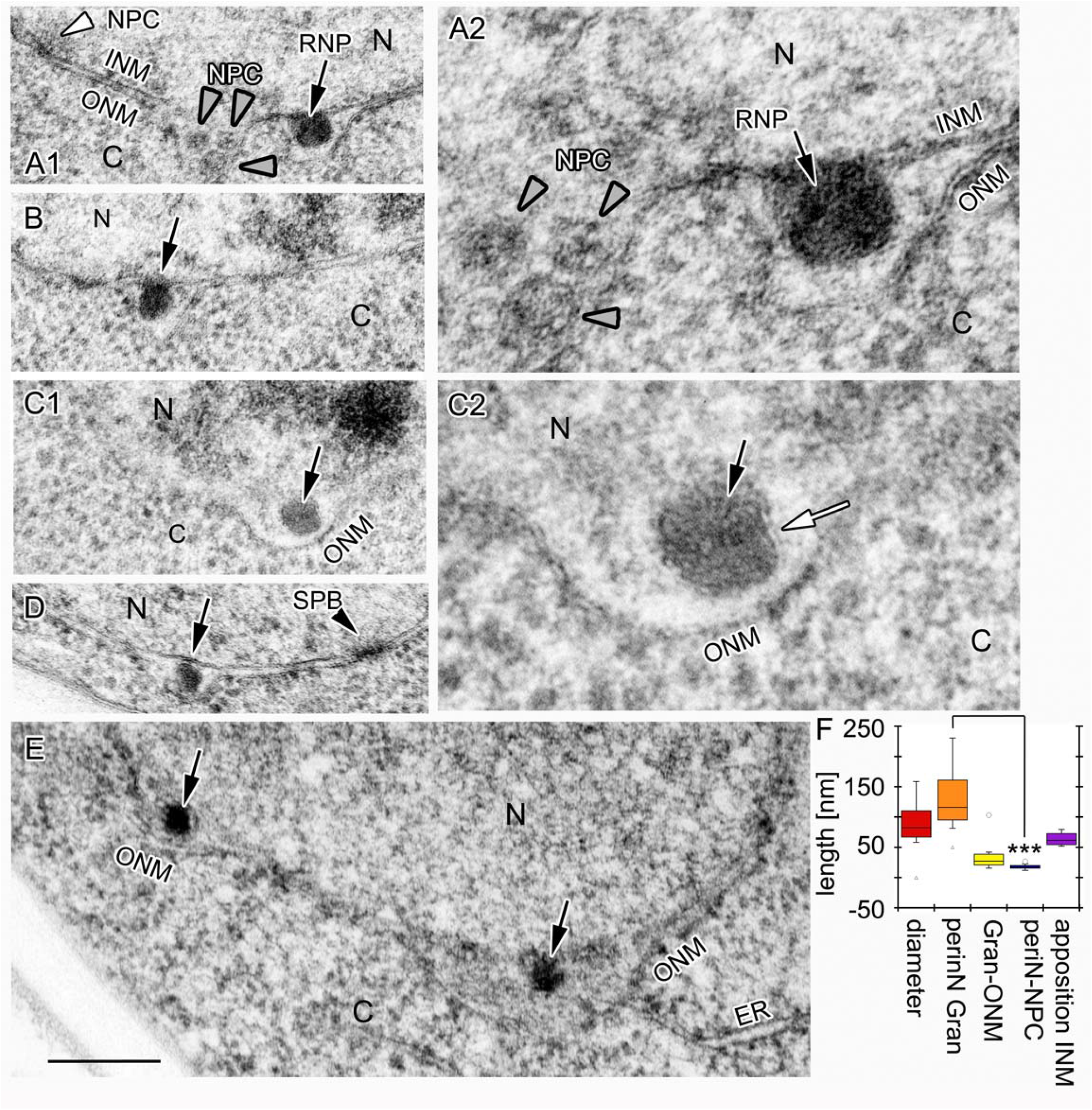
MegaRNP-like granules are observed in the perinuclear space of WT yeast. (A-E) Transmission electron micrographs of nuclei from different WT yeast cells, showing the presence of electron dense megaRNP-like granules (black arrows) present within the perinuclear space. (A) Comparison between the electron dense granules and NPCs shown at (A1) low and (A2) high magnification, as well as NPCs sectioned in a tangential (grey arrowheads) or cross-sectional (white arrowhead) plane. Note that NPCs differ both in size and shape from granules. (B) Micrograph of another megaRNP granule (black arrow) example in a different cell. (C) Micrograph of a granule (black arrow) at (C1) low and (C2) high magnification, showing that the granule at the perinuclear space is bounded by membrane (white arrow), and is separate from the ONM. (D) Comparison of a megaRNP granule (black arrow) with an spindle pole body (black arrowhead). (E) Another example of a yeast nucleus showing two megaRNP granules (black arrowheads) at the perinuclear space. (F) measured distances: diameter of the granule, measured in the orientation parallel to the nuclear envelope; perinN Gran is the size (distance) of the peri nuclear space at the site of a granule, measured at its widest point and perpendicular to the inner nuclear membrane; Gran-ONM is the size (distance) from the edge of a granule to the ONM; periN-NPC is the size of the peri nuclear space at a region of the NE adjacent to an NPC; and apposition INM is the size of the region of the granule directly apposed to the INM. N=nucleus; C=cytoplasm. Scale Bar is 500 nm for A1,B,C1,D,E and 150 nm for A2,C2.

**Figure 2.**
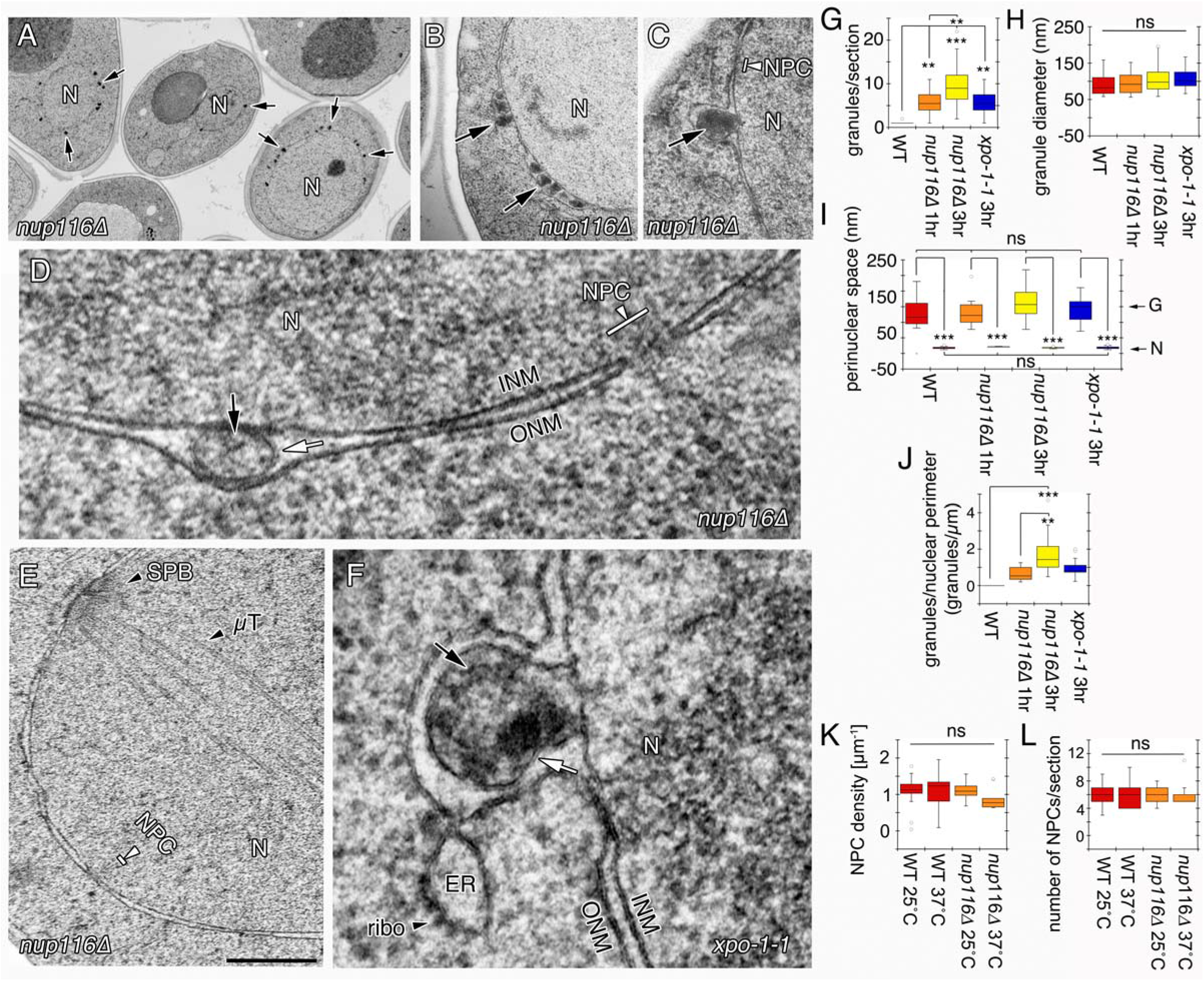
The number of megaRNP granules increases in *nup116*Δ mutants. (A) Low magnification view of *nup116*Δ mutant yeast incubated for 3 h at 37 °C, displaying many megaRNP granules (black arrows) at the NE. (B-D) higher magnification views of the granules (black arrows) showing (B) a group of granules at the perinuclear space, (C) a granule adjacent to an NPC, and (D) a granule clearly showing the surrounding membrane. (E) A view of the nucleus of a *nup116*Δ mutant cell, showing that the spindle pole body and its microtubules (μT) are normal in the mutant. (F) megaRNP granules (black arrow) similar to those observed in *nup116*Δ are also observed in *xpo1-1* mutants. White arrow points to the membrane surrounding the granule. ER=endoplasmic reticulum; N=nucleus; ribo=ribosome. Scale bar is 2.5 μm for A, 800 nm for B, E, 330 nm for C, 150 nm for D and 100 nm for F. (G-L) Quantification of granule and NPC parameters for the indicated genotypes and times: (G) number of granules per section, (H) granule diameter, (I) width of the perinuclear space, (J) granule density, (K) NPC density, and (L) number of NPC per section. Error bars represent mean ± standard error. ** p<0.01, *** p< 0.001; statistical significance was calculated by using an ANOVA with Tukey post hoc analysis.

Given the similarity in size and location of these nuclear granules to the megaRNPs that were observed in *Drosophila* (Speese et al., 2012), we sought to determine if the granules detected here in wild-type (WT) yeast might correspond to an analogous export pathway. Granules such as these previously observed in mutant yeast cells had been proposed to be artifacts caused by accumulation of material that cannot be properly transported through defective NPCs (Wente and Blobel, 1993). Similar structures were observed in strains in which NPC insertion and/or repair was impaired (Webster et al., 2014; Webster et al., 2016). To address this possibility, we compared the nuclear granules in WT cells to those from several mutant yeast strains.

### Granule frequency increases at the perinuclear space in *nup116*Δ cells

Yeast Nup116 (mammalian Nup98) is a non-essential phenylalanine-glycine-rich (FG)-Nup protein found in the central core of the NPC (Katta et al., 2014; Strambio-De-Castillia et al., 2010). Its deletion partially impairs NPC function, slows cell growth at 23 °C and becomes lethal after longer exposure at 37 °C (Wente and Blobel, 1993). Consistent with the previously published data (Wente and Blobel, 1993), we observed a substantial increase in the frequency of granules within the perinuclear space in *nup116Δ* cells after shifting the cells to the non-permissive temperature of 37 °C for 1 or 3h (Fig. 2 A-D, G, black arrows). The morphology of the granules (size, electron density, and relationship with the NE) was indistinguishable from those observed in WT yeast (Fig. 2 A-D, G,H). In addition, as in WT, the granules were enfolded by membrane (Fig. 2D; white arrow). We also observed that these granules were located at multiple sites spread out along the perinuclear space, and like in the WT, the perinuclear space at these regions was enlarged, appearing as evaginations of the ONM (Fig. 2B-D; Suppl Movie S1). When shifted to 37 °C for 3 h, 100% of the *nup116*Δ cells contained at least two perinuclear granules per TEM slice (>100 cells examined), with an average of ~ 8 perinuclear granules per section, and a range of 2-20 perinuclear granules per section (Fig. 2G). The average perinuclear granule density was 1.6/μm of nuclear perimeter (Fig. 2J). Other morphological features of the NE, such as the SPB, were not altered in the mutants (Fig. 2E).

In contrast to the previous study (Wente and Blobel, 1993), no obvious NPC-like dense structures were observed at the base of the granule or at the junction between the granule and the INM. NPC structures were frequently observed at sites outside the enlarged perinuclear space of the NE (Fig. 2C,D; white arrowheads). Importantly, both the density and the total number of NPCs were not significantly altered at 37 °C (Fig. 2K,L), even when the frequency of granules at the perinuclear space significantly increased in the *nup116*Δ mutant. In summary, several lines of evidence suggest that the observed granules are not directly derived from NPCs that became defective: (1) NPC density and number do not change with the frequency of granules, indicating that NPCs are not consumed as granules are formed; (2) the spacing of the perinuclear space around the granules is substantially larger than around the NPCs; and (3) we detect no obvious co-localization of NPCs or NPC-like density around the base of the granules. Similar conclusions were reached after observation of *ooc6/torsin* mutant-derived granules in *C. elegans* (VanGompel et al., 2015). In these mutants, NPCs become clustered in so-called “plaques”. However, when the NPCs are examined with immunoEM, it was found that while pIaques where labeled with antibodies to NPCs, the granules/blebs were not, suggesting that the accumulation of granules might push NPCs together to form the plaques. In contrast, the presence of NPC components, as observed by immunoEM, was reported in similar-looking structures in *vps4Δpom152Δ* mutant yeast cells (Webster et al., 2014). These structures were suggested to be a compartment termed the Storage of Improperly assembled Nuclear Pore Complexes (SINC), which is dedicated to the surveillance of defective NPCs in the nucleus (Webster et al., 2014). However, the distribution and morphology of the granules differ, so it is unclear if those structures truly represent the same granules described in this study, and whether they contain RNA (see below). Similarly, in TorA mutant HeLa cells, perinuclear granules displayed NPC-like immunoreactive signal at the base of the granules, and the neck of the granules abutting the INM was the same size as NPC diameters (Laudermilch et al., 2016). As yeast cells contain no identifiable Torsin homolog, it is unclear whether these are actually related structures or not.

### Granule frequency increases in *xpo1-1* mutants

To test if the accumulation of granules is specific only to mutations in NPC genes, we examined a temperature sensitive mutant of *XPO1*, which encodes the major yeast export factor CRM1 (Hodge et al., 1999; Stade et al., 1997). Large electron-dense granules, indistinguishable from those observed in *nup116Δ* mutants, were observed in the *xpo1-1* mutant after shifting to 37 °C (Fig. 2F,H). Similar to WT and *nup116Δ* cells, these granules localized within the perinuclear space (Fig.2F; black arrow), were bounded by a membrane (Fig.2F; white arrow), and had a similar average size of ~135 nm (134±6.3) (Fig. 2F,H). Unlike the *nup116*Δ cells, incubation of *xpo1-1* cells at 37 °C for 3 h resulted in granules in only ~20% of the *xpo1-1* nuclei (Fig. 2G). The granule density in *xpo1-1* was also lower, containing ~1.0 (0.97 ± 0.12) granule per μm of nuclear perimeter in positive nuclei (Fig. 2I). These differences from the *nup116*Δ granules are likely due to differences in the temperature sensitivity of this *xpo1-1* allele, not necessarily differences in NPC mutants vs. export mutants.

As disruption of the Xpo1 (Crm1) pathway leads to an increase in the frequency of granules, and Xpo1 is not an integral component of the NPC, we conclude that accumulation of these granules is not specifically caused by direct NPC structural dysfunction. In support of the model that granules do not correspond to defective NPCs, and that this is an alternative nuclear export pathway, similar granules were observed in mutants in which NE integrity or function has been compromised, including mutations in Acc1 (Schneiter et al., 1996), Apq12 (Scarcelli et al., 2007), Brr6 (Hodge et al., 2010), and Npl4 (DeHoratius and Silver, 1996).

### Granules contain mRNA transcripts

We next asked what potential cargos are contained within these granules. To examine whether these granules contain mRNA transcripts, *in situ* hybridization and TEM were combined to identify poly(A) RNAs in granules of the *nup116*Δ yeast cells using digoxigenin (dig)-labeled oligo-dT probes, followed by immunostaining of the TEM grid via gold-conjugated anti-dig secondary antibodies. As expected, gold particles were present at low levels throughout the nucleus and cytosol, identifying typical cellular poly(A) mRNA distribution (Fig. 3). Moreover, gold particles accumulated in a subset of granules in the perinuclear space (Fig. 3A-D). The number of gold particles varied from granule to granule, perhaps related to the variability of electron density observed in different granules (Figs. 1, 2). Gold particles were very rarely detected in the samples that were pre-treated with RNAse, or in the absence of the probe (Fig. 3E,F) indicating that the signal from the oligo-dT probe was specific. Not every granule showed immunoreactive signal, which could be due to fixation conditions, the export of RNAs lacking a poly(A) tail, or the use of this pathway for the export of other non-RNA materials. The presence of these gold-labeled granules in the perinuclear space supports the model that these granules contain ultra-large mRNP particles.

**Figure 3.**
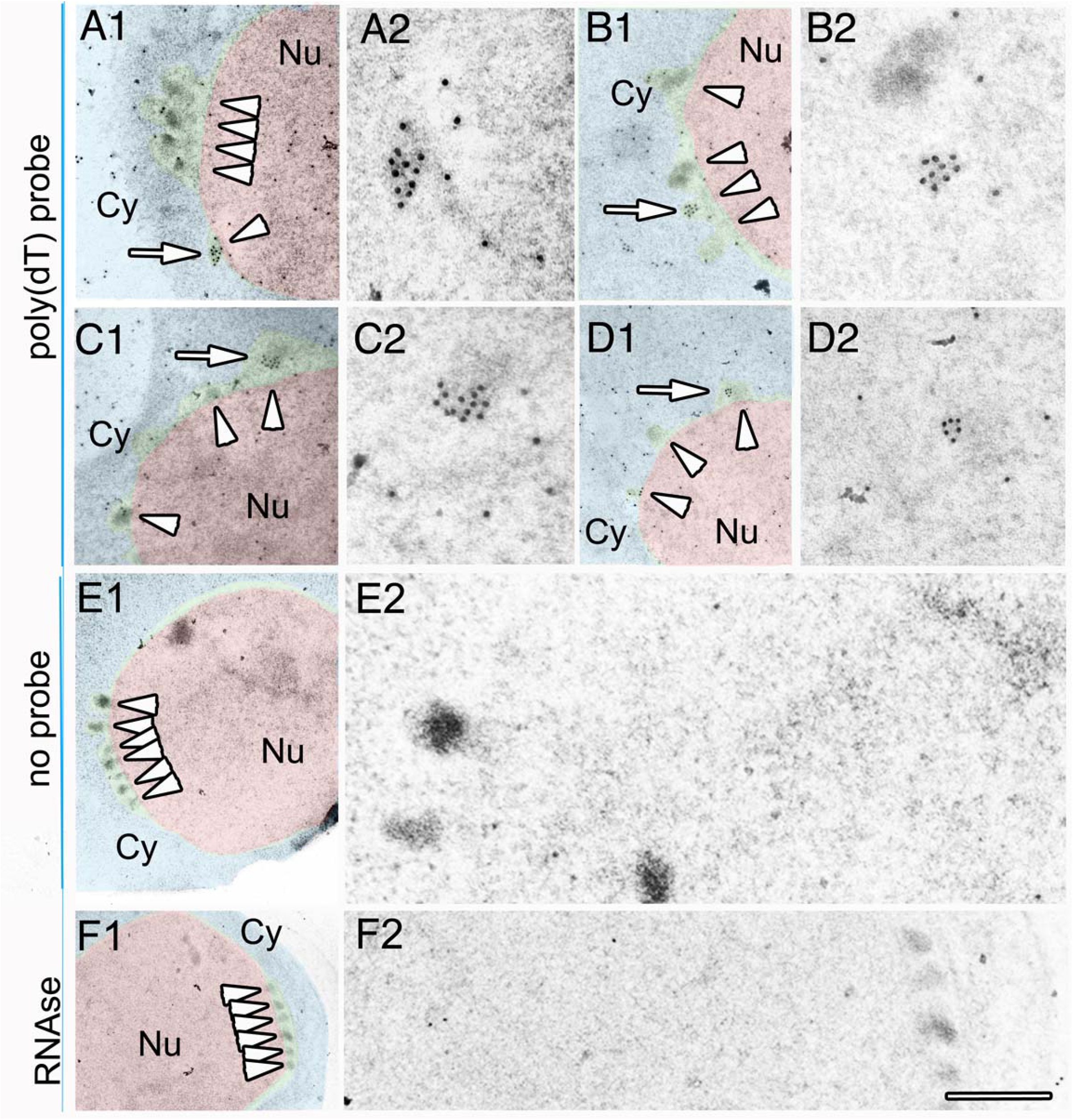
MegaRNP granules contain poly(A) RNA. (A-D) Examples of megaRNPs in *nup116*Δ cells (arrowheads) containing oligo(dT) signals detected at the EM level using 10 nm gold conjugated secondary antibodies. Labeled granules are indicated by white arrows in A1-D1, and shown at high magnification in (A2-D2). (E) megaRNP granules (arrowheads) in samples in which the oligo(dT) probe was omitted shown at (E1) low and (E2) high magnification. (F) megaRNP granules (arrowheads) in samples using a oligo(dT) probe, and treated with RNAse shown at (F1) low and (F2) high magnification. Nu=Nucleus; Cy=cytoplasm. Scale Bar is 850 nm for A1, B, C1, D1, 350 nm for A2, B2, C2, D2, F2, 300 nm for E2, and 1.5 μm for E1, F1.

### NE granules are developmentally regulated and show a polarized nuclear distribution

We examined the *nup116Δ* granules in more detail to understand when this pathway is likely to be used throughout the cell cycle. Images from TEM sections were categorized into three groups based on the presence or absence of the daughter cell (bud) and the size of the bud: no bud, small buds and large buds. Only cross-sections that displayed a continuous “neck” between the mother cell and the bud were analyzed in this study, ensuring that the cross-section of the bud was medial (e.g., Fig. 4D1, E1). This was important in order to assess the size of the bud, and thus the cell cycle stage. Given different sectioning angles, it was not possible to uniquely identify G1 cells (without buds), as a single TEM section could have missed the plane of section containing the bud. Cells with small buds were considered to be in S phase (Fig. 4D1) and cells with large buds in G2/M phase (Fig. 4E1). The average granule number in G2/M phase (~12/slice) was significantly higher than that of S phase (~6/slice) (Fig. 4A). We estimated the granule numbers of G1 cells by the calculating the total number of granules of all analyzed cells, and subtracting the granules of those in the S and G2/M phases (see Methods for details of calculation). The number of granules in G1 were estimated ~2/slice (Fig. 4A,C). Thus, granule number is positively correlated with growth stage, with more granules present during G2/M. Interestingly, the distribution of granules was asymmetric with respect to the growing bud; a larger number of granules faced the bud with the ratio (bud-facing : facing opposite to the bud) of ~2 in both S and G2/M stages (Fig. 4B,D,E). These observations showed that granule formation is upregulated during cell growth and it is polarized. The polarized distribution of granules towards the bud raise the possibility that cargos are involved in the rapid growth of the daughter cells.

**Figure 4.**
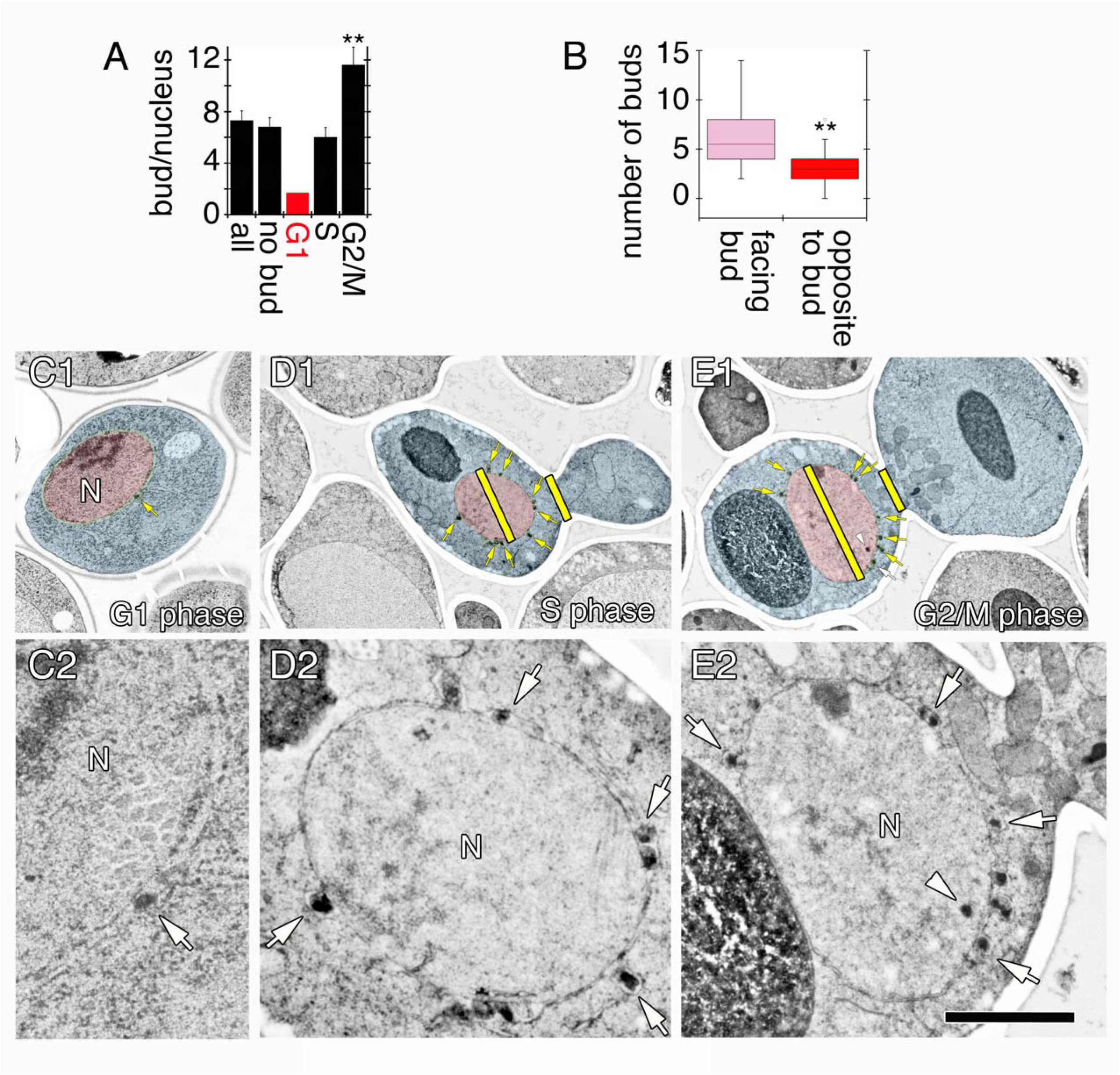
Increase in megaRNP granule frequency during the cell cycle and polarized localization of granules. (A) Quantification of the average numbers of granules per nucleus in: total nuclei: All, no bud, G1 (calculated, see Methods for details), S and G2/M phases. ** p<0.01. (B) The distribution of granules in S and G2/M phases presented as the ratio of granules in the side facing the bud to that of the side opposite to the bud. ** p<0.01. Representative images of *nup116*Δ yeast cells after being shifted to 37 °C for 3 h. (C1,2) without a bud (likely G1 phase); (D1,2) cells with small buds (S phase); and (E1,2) cells with large buds (G2/M phase). The line drawn across the middle of the nucleus separates the side of nucleus facing the bud from the side opposite to the bud. Scale bars= 3 μm (C1), 3.5 μm (D1, E1), 0.7 μm (C2), 0.8 μm (D2), 1.3 μm (E2).

### Comparison of these granules with the megaRNPs observed in *Drosophila*

The granules observed in yeast show similarities, but also some differences from the megaRNPs in *Drosophila*. Both sets of granules are substantially larger than the NPC pore, with an average diameter between 100 - 200 nm; when observed at the ultrastructural level, both are present within the perinuclear space and are surrounded by a bounding membrane derived from the INM; and both contain mRNAs, although the composition of the cargos may be distinct. Additional differences lie in the mechanisms underlying the formation of these granules. Unlike the requirement for the AAA+ ATPase Torsin in *Drosophila* and mammalian cells (Laudermilch and Schlieker, 2016), yeast contain no recognizable Torsin homolog. Likewise, the INM of yeast is not supported by a lamin-type structure (Meseroll and Cohen-Fix, 2016). Thus, the molecular details of granule invagination and scission into the perinuclear space appear to differ between species.

### Final summary

Taken together, our studies suggest that yeast contains a novel mRNA export pathway that is used under conditions in which the NPC or Xpo1 RNA export function is impaired. This pathway appears to be used at low frequency in WT cells, and is clearly not a major pathway for nuclear export, at least under laboratory conditions. We suggest that this pathway may be a more ancient conserved pathway that is inefficient compared to transport through the NPC. The observed electron dense granules are similar to those previously observed in *Drosophila* (Jokhi et al., 2013; Speese et al., 2012) and *C. elegans* mutants (VanGompel et al., 2015). It is also similar to the pathway used by Herpes family viruses as they exit the nucleus, although several of the relevant mechanistic proteins are encoded by the Herpes genome (Darlington and Moss, 1968; Hagen et al., 2015; Lee and Chen, 2010; Lye et al., 2017; Mettenleiter et al., 2013; Roller, 2008). As NPC frequency was not altered in either *nup116Δ* or *xpo1-1* mutants, this suggests that the granules are unlikely to be artifacts of defective NPC, and are likely to be intermediates along an alternative nuclear export pathway. The molecular mechanisms of this pathway do not appear to be strictly conserved from yeast to *Drosophila*, as yeast contain no recognizable TorsinA homolog and no lamins, which play roles in the upregulation of this pathway at the neuromuscular junction in flies. The underlying core machinery may be conserved, but these factors still need to be identified. It also remains to be determined under what conditions WT yeast might use this pathway, perhaps under stress or when growth is upregulated, such as initial growth of spores, a return to growth after starvation or desiccation, or after upregulation of specific cargos. It will also be interesting to further investigate the possible cargos in these granules and identify the underlying machinery for packaging and nuclear export.

## Materials and methods

### Yeast strains

The yeast strains used in this work are as follows: *nup116*Δ (SWY27; *nup116-5::HIS3 ade2-1 ura3-1 his3-11,15 trp1-1 leu2-3,112 can1-100;* (Wente and Blobel, 1993)); *xpo1-1* (*ade2-1 ura3-1 his3-11,15 trp1-1 leu2-3XPO1::LEU2* pKW457(*xpo1-1/HIS3*); (Hodge et al., 1999)); W303 (YOL2; *ade2-1 ura3-1 his3-11,15 trp1-1 leu2-3,112 can1-100;* (Wente and Blobel, 1993)). Standard methods were used for yeast media and genetic manipulations. All strains were grown at 25 °C (unless otherwise indicated) on rich media (YPD) or synthetic complete (SC) media minus the appropriate amino acid (Dunham et al., 2015).

### Transmission Electron Microscopy, Electron Tomography and immuno-EM

Electron microscopy was performed as previously described (Walther and Ziegler, 2002). In brief, for control conditions, the cells were grown to an OD of 0.5-1.0 in YPD medium at 25°C and pelleted. For heat treated samples, the cells were grown to an OD of 0.15 – 0.2 in YPD medium at 25 °C, and then shifted to 37 °C and incubated for 1 or 3 h before collection of the pellet. The fresh pellet was mixed with the same volume of 2 % low melting temperature agarose (Fisher Scientific) and subjected to high-pressure freezing (Leica EM PACT2). The frozen samples were immersed in substitution media, 2 % (w/v) osmium tetroxide, 1 % (w/v) uranyl acetate, 1 % (v/v) methanol and 5 % (v/v) water in acetone (pre cooled to −90 °C). A computer-controlled substitution apparatus (Leica EM AFS2) was used to slowly warm the samples from −90 °C to 0 °C over a period of 22 h. The samples were kept at 0 °C for 1 h, washed with water-free acetone and then followed by the stepwise embedding in epon (Electron Microscopy Sciences) mix with propylene oxide (EMS) (30% epon, 60% epon, 100% epon; each step for 1h). Resin blocks were polymerized at 55 °C for 48 h and sectioned with a diamond knife (Diatome) on a Leica Ultracut UCT ultramicrotome (Leica Microsystems). The thickness of each section was 80nm. Sections were collected on formvar coated copper grids and post-stained with 2% uranyl acetate in water and lead citrate. Images were acquired on a Phillips CM10 transmission electron microscope equipped with a Gatan CCD camera system (Erlanghsen 875) with tungsten source at 100kV, and analyzed using Gatan’s DM3 software.

For ET, resin blocks were cut in a series of 150nm-thick sections. The sections were collected onto formvar coated copper grids and post-stained with 2% uranyl acetate, followed by lead citrate. The sections were analyzed at FEI (Hillsboro, OR). Images were acquired on a Tecnai Spirit BioTWIN operating at 120kV on an FEI Eagle HS CCD camera. Tilt-series were acquired at magnifications of 30k. Each tilt-series was acquired with a range of ±70 degrees using tilt increments of 1 degree. Dual-axis acquisition was performed on both samples to improve reconstruction quality (e.g. at each location, two full ±70 degree tilt-series were acquired by tilting the specimen 90 degrees around two orthogonal axes). The reconstruction algorithm used was SIRT with 20 iterations. Data were collected by using the FEI Tomo4.0 acquisition software. The 3D reconstructions were made using the FEI Inspect 3D4.1 software and finally the visualization, segmentation and movies were created using FEI Amira 5.6.

For immunostaining samples, 0.3 % glutaraldehyde in acetone was used as substitution media after high-pressure freezing and the cells were embedded with Lowicryl K4M (EMS, 14335) and cured under UV at −30°C for 48 h, and additional 48 h at room temperature. Sections were mounted on uncoated 200-mesh nickel grids and then incubated in 20 mM Tris-HCl (pH 9.0) at 95 °C for 2 h for antigen retrieval. Grids for *in situ* hybridization controls were incubated in RNA digestion solution (100 μg/ml RNase A, 20 mM Tris-HCl (pH 7.4), 10 mM NaCl) at 37 °C for 1 h. To examine poly (A) transcripts, grids were incubated with digoxigenin-labeled oligo-dT probes in hybridization buffer (20 % formamide, 5x SSC, 50μg/ml Heparin, 0.1 % Tween-20, 0.1 mg/ml Salmon Sperm DNA, 1 mg/ml *E. coli* tRNA in nuclease-free H_2_O) at 37 °C overnight. Grids were washed with hybridization buffer 5 times (5 min each) and then incubated with blocking buffer (5 % normal goat serum (Fisher Scientific), 2 % BSA-C and 0.1 % cold water fish skin gelatin (Aurion)) at room temperature for 30 min. After wash with PBS-T (0.1% Tween-20) 5 times (5 min each), grids were incubated with primary antibody, sheep anti-digoxigenin (1:1000, Roche, #11 333 089 001) in 0.1 % BSA-C in PBS at room temperature for 1 h. The grids were washed with PBS-T for 5 times (2 min each), and then incubated with 18 nm gold-labelled Donkey anti-sheep secondary antibody (1:200, Jackson ImmunoResearch, #118799) at room temperature for 2 h. After rinsing and being completely air dried, samples were visualized with a Phillips CM10 transmission electron microscope.

### Granule frequency, size, density quantification and cell cycle analysis

Low magnification TEM pictures were taken at random; only the nuclei with definitive nuclear membranes were counted. The granule frequency was calculated from the number of granule-positive nuclei divided by the total number of nuclei that were counted. The diameter of granule size was measured using ImageJ software (NIH) using the high magnification TEM pictures of the granule-positive nuclei. For each granule, the length of the long and the short diameters were measured and the average was calculated to represent the diameter size of the granule. The granule density was calculated as the number of granules divided by the perimeter of the nucleus.

For the cell cycle analysis, based on the presence or absence of the bud (daughter cell) and the size of the bud in the low magnification of EM pictures, three groups were categorized, no bud, small bud and large bud. To ensure that the cross-section of the bud was medial, only cross-sections that displayed a continuous “neck” between the mother cell and the bud were analyzed in this study. The group of cells with small bud indicate S phase and cells with large bud indicate G2/M phase. The granule number was counted and the average (granule number per nucleus) was calculated. However, the group of cells without buds could be any grow stages because of the sectioning angle of EM samples. Based on the time of each cell cycle stage (Brewer et al., 1984), the ratio of cell number at G1, S, and G2/M phases was estimated 2:1:3, by which the number of nuclei at each stage was calculated. The granule number per nucleus (single slice) of G1 phase was calculated using the formula:

GG1 = [GT – GS*NS – GG2/M * NG2/M]/NG1

GG1, the average of granule number per nucleus at G1 phase (to be calculated)

GT, the total number of granules were counted in a set of samples (counted)

GS, the average of granule number per nucleus at S phase (calculated)

NS, the number of nuclei at S phase (calculated)

GG2/M, the average of granule number per nucleus at G2/M phase (calculated)

NG2/M, the number of nuclei at G2/M phase (calculated)

NG1, the number of nuclei at G1 phase (calculated)

For cells with bud (S or G2/M stages), the granules were counted and divided into the side facing the bud and the side opposite to the bud according to the localization. The distribution of granules in S and G2/M stages was presented as the ratio of the average of granule number in the side facing the bud to that of the side opposite to the bud.

### Statistics

Unpaired two-tailed Student’s t tests were run for comparisons of one experimental sample with its control. One-way analysis of variance was performed when comparing multiple samples, with either a Tukey (when comparing samples to each other) or Dunnett (when comparing samples to a control) post hoc tests. Results were expressed as mean ± SEM, and p < 0.05 was considered significant.

## Acknowledgments

We thank Linda Hassinger, William Hassinger, Ki Young Paek and Alexandra D’Ordine for technical assistance. Many thanks to Melissa Moore, Susan Wente, Charles Cole, Patrick Lusk and others for advice and strains. Thanks to members of the Munson and Budnik labs for critical reading of the manuscript and helpful discussions. This work was supported by a Bassick Family Worcester Foundation Award to MM, and NIH grant R01 NS063228 to VB.

## ABBREVIATIONS LIST

ET,: electron tomography;
FISH,: Fluorescence in situ hybridization;
INM,: Inner nuclear membrane;
ONM,: outer nuclear membrane;
mRNP,: messenger ribonucleoprotein;
NE,: nuclear envelope;
NPC,: nuclear pore complex;
Nup,: nucleoporin;
SPB,: spindle pole body;
TEM,: transmission electron microscopy;
WT,: wild type.

## Suppl movie-S1 Electron Tomography of granules from *nup116*Δ yeast cells

Electron tomography showing the three-dimensional structure of megaRNPs in *nup116Δ* yeast cells after being shifted to 37 °C for 1 h. The granules (red) localize in the perinuclear space, between the inner nuclear membrane (purple) and the outer nuclear membrane (blue). They contain a variety of differently sized particles (red) that are closely wrapped by inner nuclear membrane.

